# Genetic diversity in chimpanzee transcriptomics does not represent wild populations

**DOI:** 10.1101/2021.06.27.450107

**Authors:** Navya Shukla, Bobbie Shaban, Irene Gallego Romero

**Affiliations:** School of BioSciences, The University of Melbourne, Royal Parade, 3010, Parkville, Australia; Melbourne Integrative Genomics, University of Melbourne, Royal Parade, 3010, Parkville, Victoria, Australia; Centre for Stem Cell Systems, Faculty of Medicine, Dentistry and Health Sciences, The University of Melbourne, 30 Royal Parade Parkville, Victoria 3010, Australia; Center for Genomics, Evolution and Medicine, Institute of Genomics, University of Tartu, Riia 23b, 51010 Tartu, Estonia

**Keywords:** chimpanzee, genetic diversity, RNA-seq, genotyping

## Abstract

Chimpanzees (*Pan troglodytes*) are a genetically diverse species, consisting of 4 highly distinct subspecies. As humans’ closest living relative they have been a key model organism in the study of human evolution, and comparisons of human and chimpanzee transcriptomes have been widely used to characterise differences in gene expression levels that could underlie the phenotypic differences between the two species. However, the subspecies from which these transcriptomic datasets have been derived is not recorded in metadata available in the public NCBI Sequence Read Archive (SRA). Furthermore, labelling of RNA-seq samples is for the most part inconsistent across studies, and the true number of individuals from whom transcriptomic data is available is difficult to ascertain. Thus we have evaluated genetic diversity at the subspecies and individual level in 486 public RNA-seq samples available in the SRA, spanning the vast majority of public chimpanzee transcriptomic data. Using multiple population genetics approaches we find that nearly all samples (96.6%) have some degree of Western chimpanzee ancestry. At the individual donor level, we identify multiple samples that have been repeatedly analysed across different studies, and identify a total of 135 genetically distinct individuals within our data, a number that falls to 89 when we exclude likely first and second-degree relatives. Altogether, our results show that current transcriptomic data from chimpanzees is capturing low levels of genetic diversity relative to what exists in wild chimpanzee populations. These findings provide important context to current comparative transcriptomics research involving chimpanzees.

## Introduction

Chimpanzees (*Pan troglodytes*) are one of two extant members of the genus *Pan*, estimated to have diverged from the ancestral human lineage between 6 and 13 million years ago [1]. With a rich evolutionary history and high amounts of genetic diversity, they have long been used as experimental models in biological research, both as model systems and to further our understanding of human-unique traits [2–4]. After the sequencing of the chimpanzee genome showed that ~99% of sequence is common between chimpanzees and humans (ignoring genomic rearrangements), efforts have focused on identifying human-specific variation of potential functional significance [5]. Over the last 13 years, RNA sequencing (RNA-seq), a relatively low-cost and high throughput technology, has emerged as one of the leading approaches in this question, with much work having been done to characterise differences in gene expression patterns across humans, chimpanzees and other primates [6]. Transcriptomic studies have compared gene expression across organ systems, and provided chimpanzee transcriptomes for various tissues [7–9], often with a focus on the brain and neural development, in order to identify the molecular basis of cognitive and behavioural differences [10]. In parallel, induced pluripotent stem cells (iPSCs) from chimpanzees, which can be derived from existing cell lines or donor animals through minimally invasive means, are rapidly becoming established as models to study developmental intermediates and other hard-to-sample tissues [11].

The majority of this data is publicly accessible through the NCBI’s Sequence Read Archive (SRA) [12]. However, not one of the 4389 RNA-seq samples classified as Pan troglodytes in the SRA specifies the subspecies of donor individuals, suggesting that transcriptomic studies are not generally considering sample subspecies in their analyses. Yet chimpanzees show population stratification greater than all other lineages of great apes [1,13,14]. They can be split into two monophyletic clades, each containing two distinct subspecies [1,15,16]. The first clade consists of the closely related Central (Pan troglodytes troglodytes) and Eastern chimpanzees (Pan troglodytes schweinfurthii), found in equatorial Africa, and the second clade is comprised of the Western chimpanzees (Pan troglodytes verus), found in upper Guinea and the recently designated Nigeria-Cameroon (*Pan troglodytes ellioti*) chimpanzees from the Gulf of Guinea [1,17,18]. Each subspecies has had a long separate history, and has been exposed to a wide range of ecological variation, such as incidence of pathogens, imposing different selective pressure [19], although evidence also points to considerable gene flow between subpopulations [14,20]. Pairwise F_ST_ between Central and Eastern chimpanzees is estimated to be 0.09, similar to values seen between different populations of humans, but F_ST_ between Western and Central chimpanzees is 0.29 and that between Western and Eastern is 0.32 – significantly higher values than typical human estimates [13]. There is also extensive diversity within individual subspecies. For example, looking at a noncoding locus on the X chromosome shared between humans and chimpanzees, the mean pairwise sequence difference among all humans is 0.037%, while among Central chimpanzees alone it is 0.18% [21].

In transcriptomic studies of chimpanzees, tissue samples are often collected *post mortem* from captive individuals, following death from unrelated causes. However, it is unclear how representative of the diversity in wild chimpanzees these small captive populations are. A genetic survey looking at European zoos and research institutes showed that the majority of individuals (70%) had some Western ancestry [22]. A similar survey of American chimpanzees also produced highly skewed results, with 95% of studied individuals being of Western descent [23]. While collections strategies in studies often aim to be random, sampling from these reservoirs with depleted genetic diversity suggests there is a strong likelihood that transcriptomic studies are not sampling much of the intra and inter subspecies diversity found in chimpanzees. Here we consider that possibility, generating genotype data from publicly available chimpanzee RNA-seq samples to investigate the state of genetic diversity in current transcriptomic research involving chimpanzees through a population genetic lens.

## Results

### Initial sample selection and genotyping

As of the 19th of January 2021, the NCBI SRA database contained 4389 RNA-seq samples with taxa label Pan troglodytes. Following stringent filtering steps (see methods), we retained 486 BioSamples from 40 different BioProjects (collections of BioSamples from a single initiative in NCBI; Supplementary Table 1, Supplementary Table 2) and 20 different tissue or cell types. We note that in the SRA BioSample is an ambiguous term that covers different experimental contexts and designs. While in one study different BioSamples can refer to genetically distinct individuals, in another they may indicate different tissue sections or different experimental treatments applied to the same individual. For example, PRJNA299472 (S7, [24]) has 72 BioSamples — these correspond to 18 different sections from the prefrontal cortex of only 4 different individuals. In the following we therefore distinguish replicate samples, which are BioSamples from the same donor animal and the same study, and *duplicate* samples, which are BioSamples from the same donor animal across different studies, defined on the basis of publicly available metadata either in SRA or the associated publication.

We successfully genotyped 6,943,957 SNPs in these 486 samples, 54,706 of which had ≤ 5% missingness. An initial UPGMA tree (Supplementary Figure 1, see methods) identified a clade of 18 samples with consistently large (≥ 0.1) Identity-by-State (IBS) distances from the rest of the samples, which we reasoned may represent sample swaps or human contamination. Examination of mitochondrial reads from these samples suggested in all cases that they were not of chimpanzee origin (Supplementary Table 3), and we thus excluded them from any further analyses. Our final dataset therefore contained 468 samples.

### Population structure and genetic ancestry of transcriptome samples

To identify the ancestry of the RNA-seq samples, we merged our genotype data with 28,559,256 SNPs genotyped in 59 wild-born chimpanzees through high coverage whole-genome sequencing [14]. Since we were merging RNA-seq-derived variants with whole genome variants, we excluded all SNPs with missingness ≥ 5% (we note that our results are robust to this choice of threshold, see Supplementary figure 2). This led to a merged dataset containing only 11,679 SNPs across all 527 samples, which we confirmed was sufficient to capture subspecies differences through principal component analysis (PCA) of the wild-born samples alone (Figure 1A, 1B). This number of SNPs was sufficient to clearly differentiate Western and Nigeria-Cameroon chimpanzees from Central and Eastern chimpanzees on PC1 and 2, and to further separate Central and Eastern chimpanzees from each other on PC3, recapitulating trends observed in the WGS dataset.

**Figure 1:**
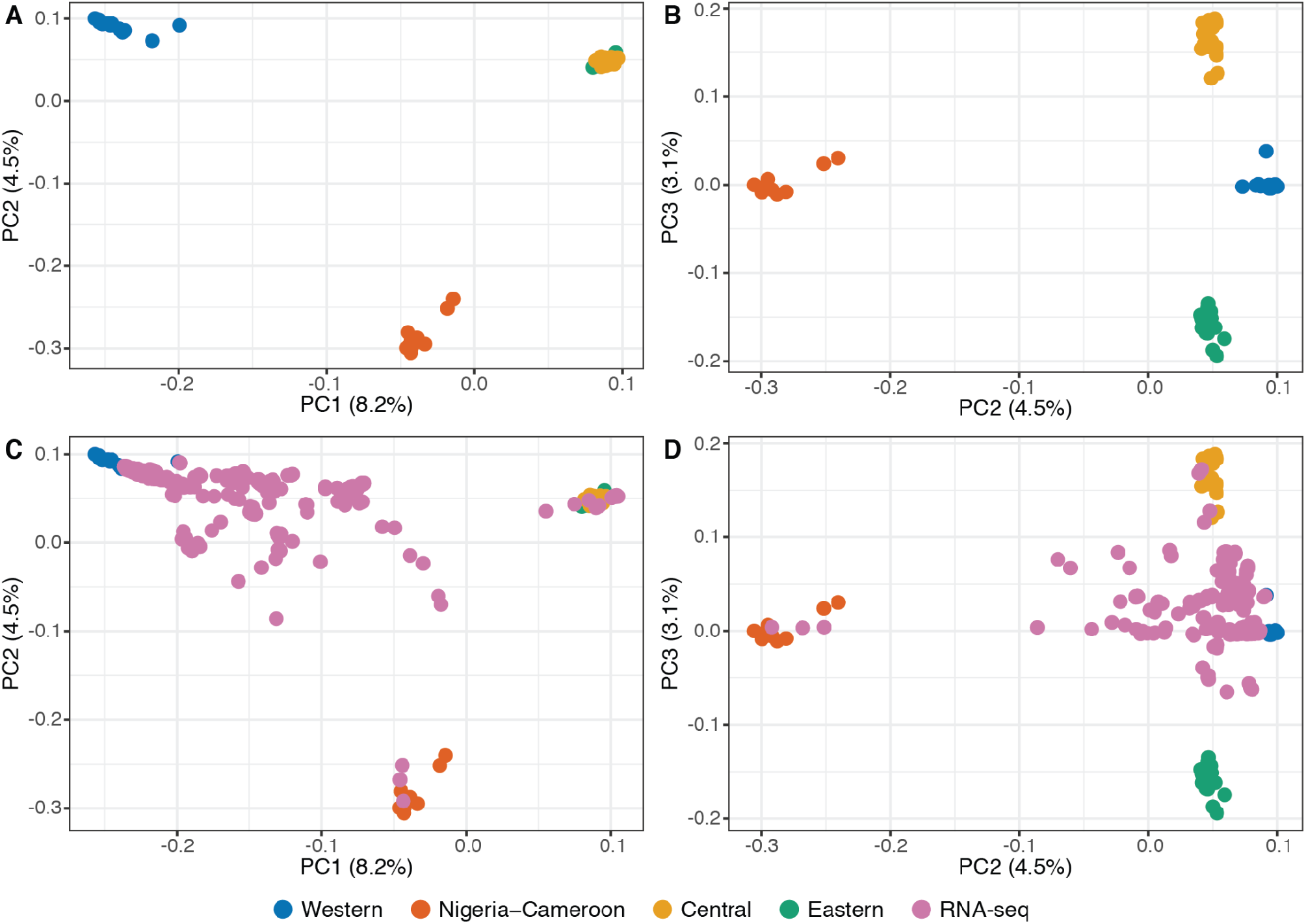
Principal component analysis of 527 chimpanzee samples with genotype data. **A, B.** PC1-PC2 and PC2-PC3 of 59 wild-born samples of known ancestry [14] using 11,679 SNPs genotyped across the entire dataset. **C, D.** Projection of 468 chimpanzee RNA-seq samples of unknown ancestry onto the PCA of wild-born samples. Colors indicate the four distinct chimpanzee subspecies and samples of unknown ancestry from public RNA-seq datasets.

We then projected the 468 RNA-seq samples onto this PCA (Figure 1C, 1D). A substantial fraction of RNA-seq samples cluster with Western chimpanzees, and a further 3 individuals fall within the Nigeria-Cameroon chimpanzee cluster. However, the majority of samples fall outside the four subspecies clusters, suggestive of mixed ancestries and of the presence of substantial levels of admixture in our dataset. Examination of PC2 and PC3 suggested that there are 4 samples of possible unadmixed Central origin in the RNA-seq data, and none of unadmixed Eastern origin.

To refine these ancestry inferences, we then performed a supervised ADMIXTURE analysis with the RNA-seq samples, using the 59 known individuals as a reference (Figure 2). The results of this analysis broadly recapitulated trends observed in the PCA, including an outsized amount of Western ancestry within samples. In determining the ancestry composition of a sample, we consider it to be associated with a particular subspecies if that subspecies contributes ≥ 6.25% to a sample’s genome, equivalent to having one un-admixed great-great grandparent from that subspecies. Using this definition, 452 of the 468 samples had some Western ancestry, and 287 were exclusively of Western ancestry (Figure 2, Table 1). The average proportion of Western ancestry among these 452 samples, including admixed individuals is 83.4%.

**Figure 2:**
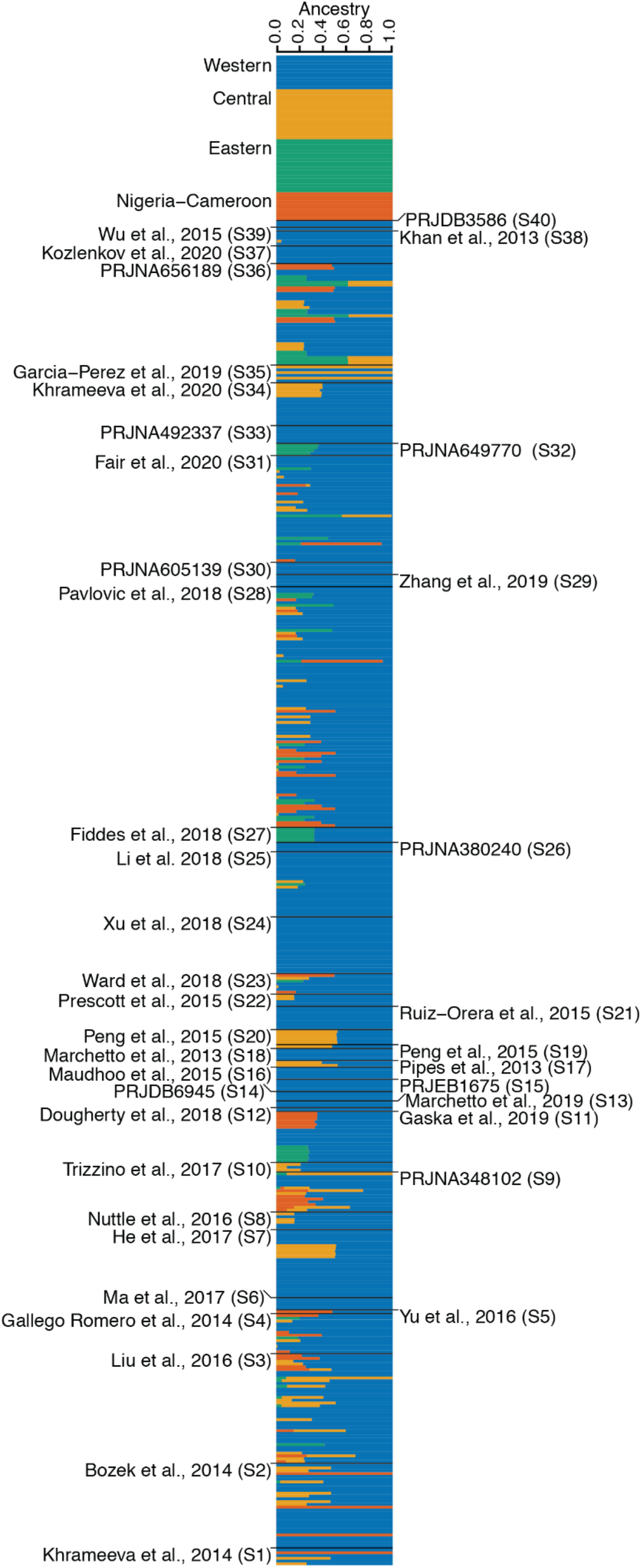
Supervised ADMIXTURE on 468 chimpanzee RNA-seq samples of unknown ancestry. Samples from [14] are shown at the top; internal study nomenclature is given in parenthesis for the 40 RNA-seq datasets with samples of unknown ancestry; full references are available in Supplementary Table 1. Within a study, samples are ordered by SRA identifier.

**Table 1:** Summary of ADMIXTURE results:

|  | Western | Nigeria-Cameroon | Central | Eastern |
| --- | --- | --- | --- | --- |
| Number of samples with at least 6.25% inferred ancestry: | 452 | 59 | 90 | 51 |
| Average inferred ancestry (%) | 87.7 | 37.7 | 35.6 | 32.6 |
| Max ancestry (%) | 99.9 | 99.9 | 99.9 | 62.2 |

Using the same criteria only 51 samples showed evidence of Eastern ancestry, with the average proportion being 32.6% and with no sample being assigned exclusively to this subspecies. In the 59 samples with some Nigeria-Cameroon ancestry this value is 37.7%, and in the 90 samples with Central ancestry it is 33.8%. As suggested by our PCA, 3 samples are predicted to be of entirely Central ancestry, and 4 are exclusively of Nigeria-Cameroon ancestry. The remaining 174 samples show substantial contributions from more than one ancestry, evidence of recent hybridisation between subspecies; 165 of these contain a substantial amount of Western ancestry, and 10 samples show substantial contributions from three of the subspecies, although none carry ancestry from all four. Most samples are either fully Western or likely had a Western parent or grandparent, making this the clear predominant ancestry source in functional genomics chimpanzee data. Reassuringly, with few exceptions, which we discuss below in more detail, both known replicates and duplicates showed consistent predictions across BioSamples and BioProjects, suggesting an overall low number of sample misclassifications/swaps in our dataset.

### Genetically distinct individuals represented in chimpanzee RNA-seq datasets

To begin examining how many different chimpanzees were represented within the 468 samples in our data, we calculated pairwise IBS distances between all samples and built a UPGMA dendogram (Supplementary Figure 3). As expected, known replicate samples clustered together. The only exception to this was a kidney sample from individual S2-c5-M-38y (naming convention for samples is “Study ID - original sample ID - sex - age;” not all of this information is available for all samples), which failed to cluster with three other replicates from the same individual. This could be a case of mislabelling, with the kidney potentially belonging to another chimpanzee. Although 296 of samples were of either neural/brain (167) or cardiac/muscle (129) origin, we observed no tissue-of-origin impact on sample clustering. Instead, different tissue samples from the same individual robustly clustered together, both within and between studies, confirming that UPGMA clustering could be used as a means of determining sample identity.

As a second layer of analysis and to further validate our replicate groupings, we calculated pairwise relatedness and identity-by-state across all 468 samples in the data using Somalier [25], a tool designed to identify sample swaps across large high-throughput sequencing datasets (Figure 3A). We then asked how many of the 530 known replicate pairs (Supplementary Table 2) had a relatedness estimate ≥ 0.5, the recommended threshold for identifying identical samples from RNA-seq data. Only 4 pairs failed to meet this threshold. Three pairs included the kidney sample from S2-c5-M-38y described above (Supplementary Figure 3); pairwise relatedness of this sample with all other S2-c5-M-38y replicates was < −1 (Figure 3A). The final pair had a relatedness value of 0.489 and included a muscle and a brain sample from the same replicate set. On the whole, relatedness estimates within sets of replicates were high, suggesting genotype sparsity and differences in tissue-of-origin did not broadly hinder our ability to identify identical samples. Excluding the three clearly unrelated pairs described above, mean relatedness amongst pairs within a replicate set was 0.884 (sd = 0.085), whereas mean pairwise relatedness in the whole dataset was −0.211.

**Figure 3:**
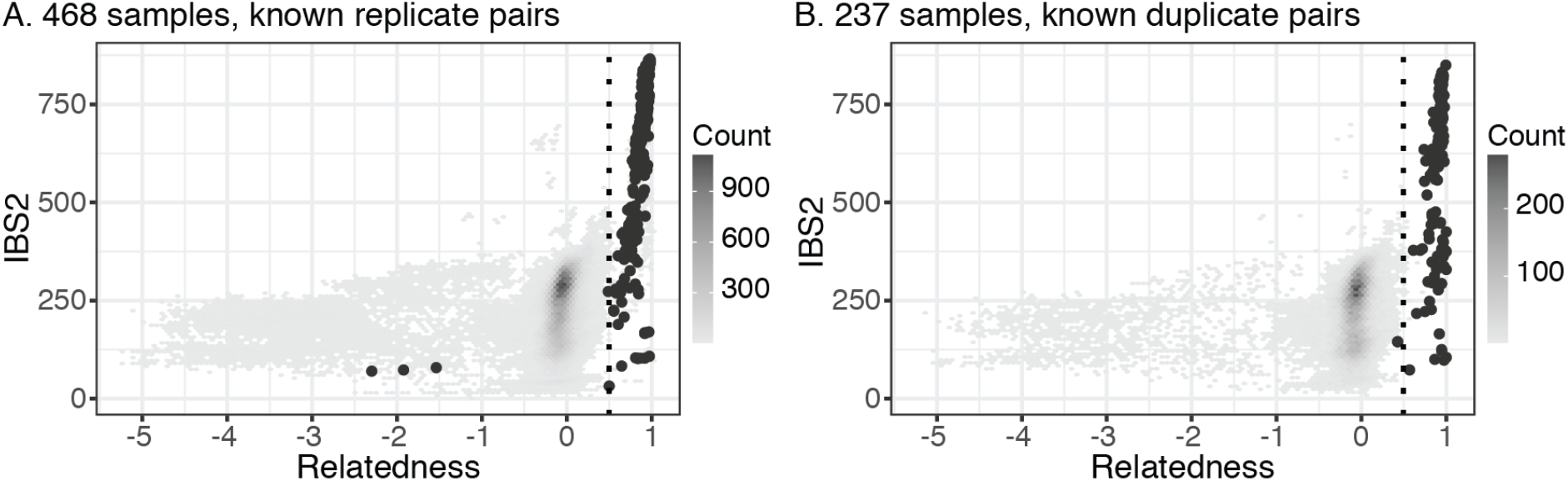
Relationships and cryptic relatedness across samples. **A.** Pairwise IBS2 and relatedness, as estimated by [25], in the unfiltered set of 468 samples. Known replicate pairs are coloured dark grey. **B.** as A, but only considering 237 non-replicate samples. Known duplicate pairs are coloured dark grey.

An additional 801 sample pairs (involving a total of 351 samples) not belonging to a replicate set also had relatedness ≥ 0.5. As the 468 samples used in these analyses included both known duplicate samples and likely given the relatively small captive chimpanzee population, close relatives, which Somalier is not designed to detect, this finding was not surprising. On the whole, these analyses suggested that we are well-powered to determine whether sets of replicate samples in our data are actually identical or not.

Given this, we then filtered our dataset to retain only one sample from each replicate set (see methods), resulting in a set of 237 samples. An UPGMA dendogram (Supplementary Figure 4) of these revealed 71 sets of duplicated samples across studies, all of which met at least one of the following criteria: clustering together with terminal node edge length < 0.006, or clustering together with node length ≥ 0.006 but supporting metadata (age, sex) in common; only two clusters (5 and 9) fell under the latter. While metadata support for most of these duplicate sets was strong, in some cases available information was scant or incongruous (e.g., Cluster 59, where sampling ages differ by 9 years despite both samples being from brain tissue likely collected *post mortem*).

We then repeated the Somalier analysis with the set of 237 samples (Figure 3B). Average pairwise relatedness between the 168 sample pairs from the same cluster was 0.900 (sd = 0.089). However, 2 pairs, both part of Cluster 69 (Supplementary Figure 4) had relatedness < 0.500. Examination of these pairs revealed that they had S37-BooBoo-M-28y in common; relatedness is 0.427 between S37-BooBoo-M-28y and two other samples in Cluster 69, and 0.803 between S37-BooBoo-M-28y and S3-c25-M-23y. In turn, S3-c25-M-23y had a pairwise relatedness of 0.944 with the other two samples in the cluster. Using combined evidence, we still assigned S37-BooBoo-M-28y to Cluster 69. The mislabelled kidney sample from S2-c5-M-38y formed a cluster with S3-c16-F-6132 (Cluster 57, pairwise relatedness 0.750), a brain sample from the Southwest National Primate Research Center (SNPRC) with no additional replicates.

We also noticed some likely sample swaps, where known duplicate samples failed to cluster together. For instance, 4 samples from S23 appeared to be mislabelled amongst themselves (Supplementary Figures 3 and 4, Clusters 7, 12, 24, and 71; the authors of the study [26]confirmed the sample swap occurred at the time of data upload to NBCI and has since been corrected; pers. comm M. Ward to IGR), while S18−PR01209−M−2y clustered with cell line PR00818 (Cluster 33) rather than with other instances of PR01209 (Cluster 28). Cluster 14 contains 3 samples with different names and metadata, S27-S008919-F-10y, S31-724-F-15y and S28-4933-F-6y, all with pairwise relatedness > 0.9. Searching in Cellosaurus [27] revealed that S008919 is a cell line that was part of the Yerkes National Primate Research Center collection available from the Coriell Institute, and which according to a partial Yerkes pedigree shared with us by B. Pavlovic, B. Fair, and Y. Gilad was established from a Yerkes chimpanzee with internal sample ID 724, confirming they are the same animal. Conversely, relatedness between S28-4933-F-6y and S11-S4933-F-6y, which should be the same individual, was 0.206. In addition, relatedness was high between S28-4933-F-6y and S28-495-M-NA (0.505) or S31-495-M-NA (0.517), which are duplicate instances of Amos, the father of S008919. We thus concluded that S28-4933-F-6y is mislabelled, and actually derived from S008919.

### Identifying cryptic relatives in public chimpanzee RNA-seq datasets

In our Somalier analyses 18 pairs of samples not assigned to the same duplicate cluster had relatedness > 0.5 (Supplementary Table 4, Supplementary Figure 5), again hinting at the presence of cryptic relative pairs in the data. Thus we filtered our data to a final set of 135 genetically unique individuals (see methods), and computed pairwise identity-by-descent (IBD) metrics for these individuals. We first analysed only the 38 samples from S31, which contains 8 known pairs of first-degree (all parent/offspring) relatives [28]. All 8 of the known pairs had IDB Z1 > 0.60, and 7 had IBD Z1 > 0.70. In addition, we were aware of 5 pairs of known second degree relatives; IBD Z1 scores for these pairs ranged from 0.19 to 0.60 (Figure 4A). We used these results to define conservative IBD Z1 and Z2 thresholds that allowed us to identify possible additional relative pairs in the rest of our data.

**Figure 4:**
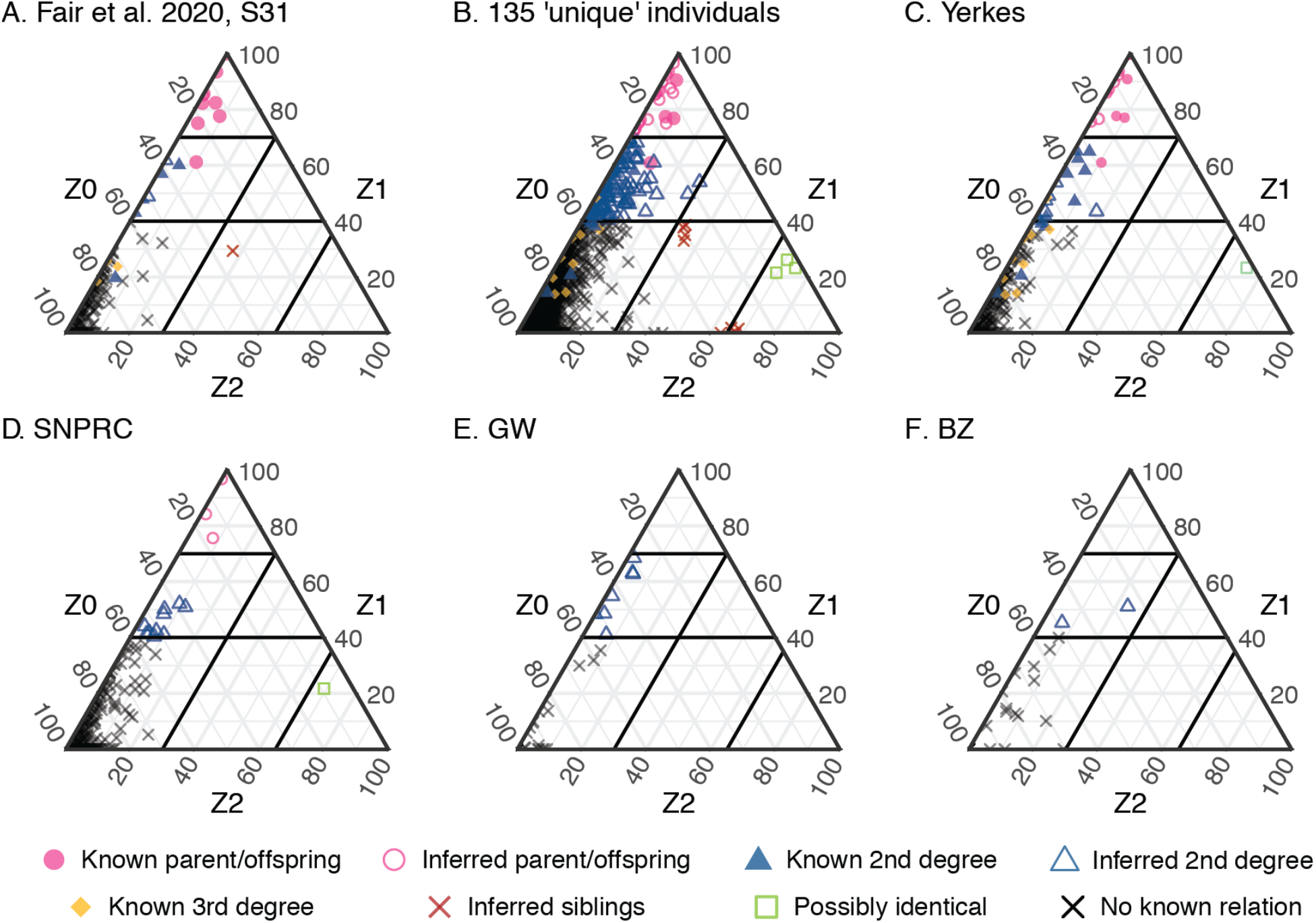
Predicted relatedness amongst 135 chimpanzee samples in public RNA-seq data. Ternary diagrams showing pairwise IBD estimates across different data subsets. Thresholds for defining different relationships between pairs are indicated by black lines on the diagram. Yerkes: Yerkes National Primate Research Center; SNPRC: Southwest National Primate Research Center; GW: George Washington University National Chimpanzee Brain Resource; BZ: Burger’s Zoo in Arnhem.

We applied the same approach to all research centers or zoos with more than 5 samples, as well as to the entire dataset (Figure 4B-F, Table 2, Supplementary Table 5). We identified only 27 pairs of first-degree relatives across the entire data set, but 146 likely second-degree relative pairs, again suggesting that the overall captive chimpanzee population is small. Of note, 97 of the likely second-degree relative pairs involved individuals sampled in different sites, highlighting connections between them. Within the Yerkes dataset (Figure 4C) we identified a pair of samples with IBD Z2 0.7455, which we predicted to be derived from the same donor individual. This assignment is partially supported by incomplete metadata for the two individuals; there are two additional cases in the full dataset (Figure 4B; IBD Z2 > 0.65 in all three cases). Below we have retained both members of all 3 pairs, but these observations suggest that the true number of genetically unique individuals in our dataset is between 135 and 132.

**Table 2.** Relatedness summary across sampling sites:

|  | All samples<br>(n = 135) | Fair et al<br>(n = 38) | Yerkes<br>(n = 36) | SNPRC<br>(n = 36) | GW<br>(n = 12) | BZ<br>(n = 9) |
| --- | --- | --- | --- | --- | --- | --- |
| Possibly identical pairs | 3 | 0 | 1 | 1 | 0 | 0 |
| Parent/offspring pairs | 27 | 9 | 14 | 3 | 0 | 0 |
| Second degree pairs | 146 | 8 | 22 | 10 | 7 | 2 |
| Full siblings pairs | 9 | 1 | 0 | 0 | 0 | 0 |

**Table 3.** SNPs preserved at missingness thresholds of 1%, 2% and 5%, for both RNA-seq data merged to reference SNPs and unmerged RNA-seq data:

| Missingness | Merged Data | Unmerged Data |
| --- | --- | --- |
| no filter | 29,968,153 | 3,081,983 |
| 10% | 18,679 | 87,196 |
| 5% | 11,659 | 54,706 |
| 2% | 5814 | 27,761 |
| 1% | 3545 | 15,494 |

Our analyses also readily identified four clear full sibling pairs, visible in the centre of Figure 4B, in the full dataset. Three of these involved the same individual, S28-462-F-42y, and three other animals, which should therefore also be full siblings with one another. Instead, these additional pairs involving these 3 chimpanzees (S3-c37-M-35y, S31-4×0354-M-21y and S36-4×0421-F-NA; Z2 between the latter two is high enough to suggest they might be the same individual) consistently showed unexpectedly high IBD Z2 values (> 0.60) and a nearly complete lack of differing heterozygous sites (IBD Z1 < 0.018 in all cases), and are visible in the bottom right of Figure 4B; a fifth chimpanzee, S35-CH114-NA-NA also exhibited high IBD Z2 values with two of the animals in this group. Intriguingly, these individuals were sampled across a variety of centers and tissues, with no obvious links between them. Despite the lack of known full sibling pairs to guide our inferences, we cautiously labelled these individuals as siblings, while acknowledging unresolved complexity in their relationships.

### Identifying unique and unrelated individuals in transcriptomic datasets

Our analyses identified between 132 and 135 genetically unique individuals in our dataset; a simple UPGMA dendogram of all 135 is presented in Figure 5. As may be expected, we observed significant reuse of samples across studies (mean number of studies an individual appears in = 1.7), with the two most frequent individuals in our final dataset being cell lines: PR00818 appeared in 9 BioProjects and C3649 in 5. A further 53 samples appeared twice, 12 thrice and 4 four times. On average, 1.35 tissues have been studied per animal, with the brain and the heart being the most commonly sequenced ones—across 58 different animals in the case of brain or neural tissue and 60 in that of cardiac or muscle tissue. Notably, these are broadly distinct sets of individuals; animals studied for their brain tended to not be considered much further (mean number of tissues sequenced = 1.29), and the same is true of cardiac samples (mean number of tissues sequenced = 1.13).

**Figure 5:**
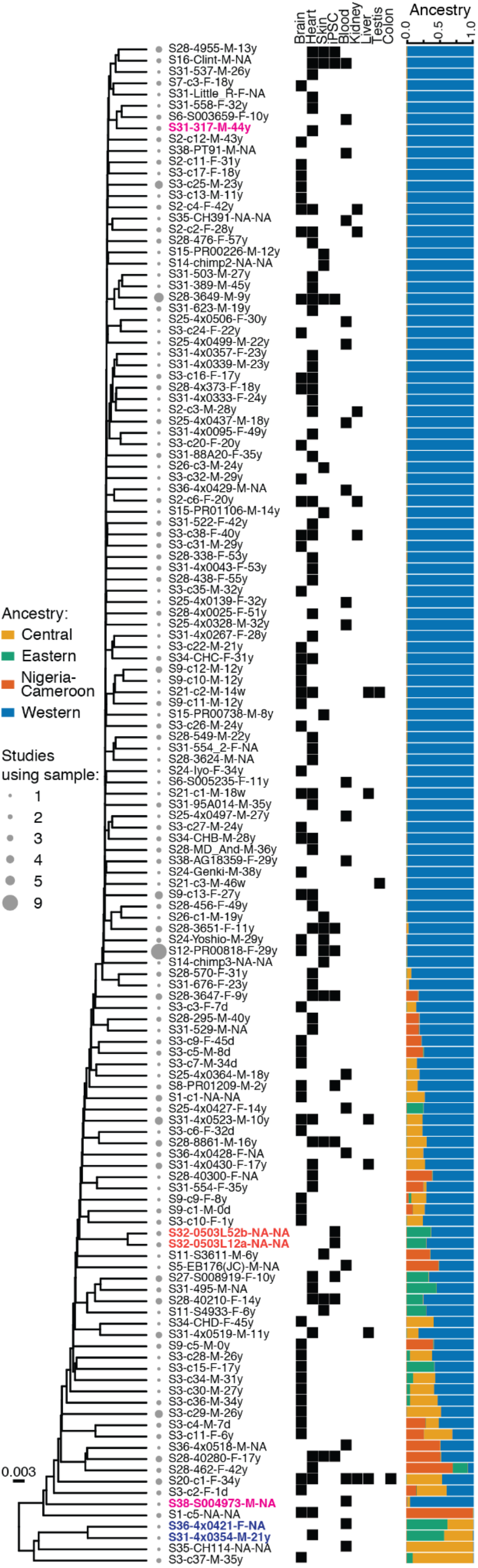
135 genetically distinct chimpanzee individuals in public RNA-seq datasets. UPGMA dendogram of 135 samples predicted to be genetically distinct in our dataset. The three sample pairs predicted to be closely related by relatedness and IBD analyses are indicated by coloured taxon labels. Filled boxes indicate tissues sampled across studies from each individual. Ancestry estimates are as in Figure 2.

As our analyses above demonstrated, these individuals are not unrelated, nor do they represent a random sample of the genetic diversity of the wild chimpanzee population. Beyond the 3 sample pairs that appeared to be genetically identical, 72 individuals had at least one second-degree relationship with another individual in the data set, and 36 were part of at least one parent/offspring or sibling pair; a single individual was part of 4. On average any individual had at least 2.09 2nd degree relatives in the dataset, and 7 had over 10; in the most extreme instance we inferred 19 second-degree relationships involving S3-c34-M-31y. We iteratively excluded individuals with the most relationships until no first or second-degree relative pairs remained in the dataset; this subset contained between 86 and 89 individuals, depending on which relative we removed from certain pairs.

Furthermore, of the 135 individuals, 85 (63%) were entirely of Western ancestry (>93.75% as predicted by ADMIXTURE), and 130 out of 135 (96%) have significant (>6.25%) Western chimpanzee ancestry. There were 48 hybrid individuals, 42 of which are of two different ancestries and 6 of three different ancestries. The final dataset contained only 1 Central chimpanzee, 1 Nigeria-Cameroon chimpanzee, and no Eastern chimpanzees at all.

Finally, we considered the subset of samples from induced pluripotent stem cells (iPSCs) or iPSC-derived cell types (Figure 6). As iPSCs can be indefinitely maintained and experimentally perturbed, and, upon directed differentiation, give access to a large number of otherwise unobservable tissue types and cell states, they have recently become established as a model system in comparative functional genomics studies [11]. However, because they are time-consuming to establish, only 15 (possibly 14, depending on how we resolve the high similarity between S32-0503L12-NA-NA and S32-0503L52-NA-NA) have been established to date; each of them has been included in an average of 2.6 studies.

**Figure 6:**
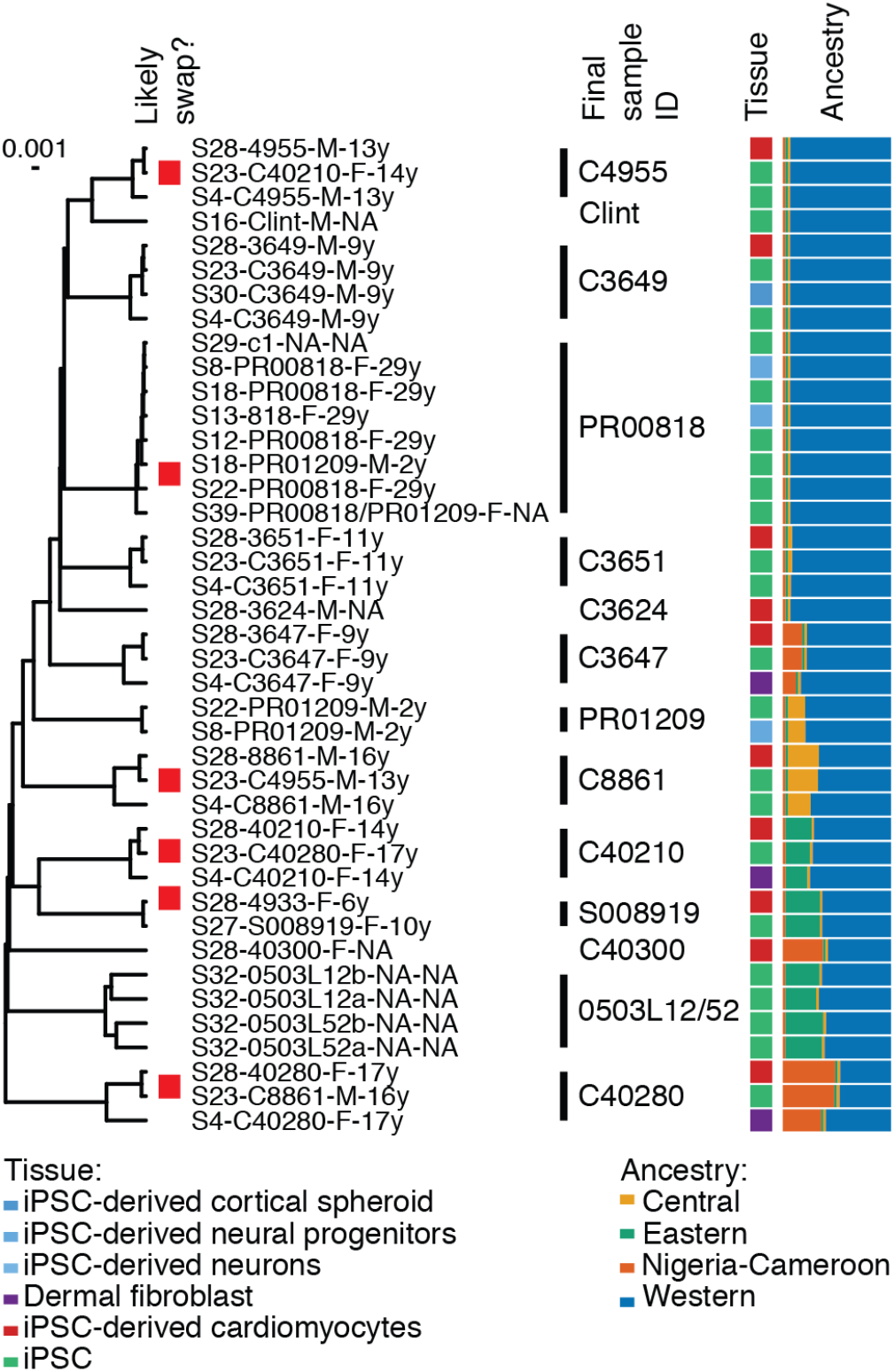
Genetic diversity and sample swaps amongst iPSC and iPSC-associated samples. Genetic diversity, tissue type and relationships amongst the 15/14 chimpanzee iPSC lines in our dataset. Ancestry estimates are as in Figures 2 and 5. Samples that appear to be mislabelled are indicated by the red box next to individual sample IDs, multiple samples predicted to be from the same genetically unique individual are marked in the final ID column.

Of the 15 different individuals in our data with iPSC data, 6 are entirely Western and the rest are two-ancestry hybrids with at least one Western parent, again highlighting the difficulty of successfully capturing existing chimpanzee genetic diversity in functional genomics resources. Additionally, our IBD analyses suggested that C40210 and C8861 are second degree relatives, as are C40210 and S008919. Notably, all but one of the sample swaps we described above involve iPSCs or iPSC-derived samples, emphasising the increased complexity of cell culture relative to observational data generation from frozen tissues [29].

## Discussion

Comparisons with chimpanzees are essential for understanding human evolution and the genetic basis of human-specific traits. But chimpanzees are a critically endangered species, and increasing their captive population is unethical, and indeed illegal in many countries. Samples are therefore strictly limited to those from animals currently living in captivity, and to *post mortem* tissue and cell line collections. These populations were established decades ago, sometimes with little regard to subspecies differences, and have not always been managed to maximise genetic diversity. Breeding practices in captive colonies, especially prior to the establishment of genetically guided breeding programs, have produced hybrids with highly variable ancestry components and this lack of specificity makes them a poor resource [22].

Our meta-analysis of chimpanzee individuals in transcriptomics provides important information for contextualizing comparative research done so far, especially as many studies do not include more distantly related outgroups. We consistently find that both in the number of samples and proportion of ancestry there is an overwhelming bias towards Western chimpanzees (*Pan troglodytes verus*) in public datasets. This is particularly noteworthy as, of the four extant chimpanzee subspecies, Western chimpanzees have the smallest effective population size, and show the highest amount of genetic drift [1,20]. All 40 studies we considered included at least one unadmixed individual of Western ancestry. In contrast, only two studies sampled a Nigeria-Cameroon chimpanzee (as we later discovered, this was the same individual); and only S35 [30] sampled a purely Central chimpanzee. None of the studies considered subspecies when sampling.

We also observe ample evidence of cryptic relationships, further pointing towards a depletion of genetic diversity. We are able to identify a total of 135 (possibly 132) unique individuals and a maximum of 89 unrelated individuals within our initial set of 468 chimpanzee RNA-seq samples. At least 82 samples have a first or second-degree relationship with another sample in the dataset. As expected from captive-bred individuals, some of these relationships are between samples from the same source. However, we see a significant number of cryptic relationships between institutes. This means that even sampling from multiple sources or from tissue repositories like GW could include some cryptically related samples.

In addition, many individuals in our data have been sampled more than once. The least concerning scenario of repeat sampling is sequencing of different tissues from the same donor—but only 29 individuals have transcriptomic data from more than one tissue type. If we consider the 58 individuals with neural RNA-seq data, 37 appear in more than one study, and 21 of those only in the context of neural samples across different studies. As half of these samples are derived from the National Chimpanzee Brain Resource at George Washington University, a *post mortem* brain bank, it is likely that these studies are repeatedly sequencing the same brain sample. Similarly, of the 60 heart/muscle samples, 40 have been used more than once, 17 only for heart/muscle purposes - all 17 have been used by both S31 [28] and S28 [31], which were conducted by the same research group using partly overlapping collections of individuals/cell lines. This finding has implications about how much unique transcriptomic information is actually available.

Unlike with human studies, however, more diverse sampling of chimpanzees is largely not possible. The only currently viable solution is deliberate sampling of captive donors of specific subspecies, and the expansion of existing iPSC collections. Unfortunately, genetic surveys of captive chimpanzees in zoos and research institutes worldwide consistently find that over 75% of captive individuals are of solely Western ancestry [22,23,32], and indeed, they are the only subspecies for which a genetically-guided breeding program has been established, by the European Association of Zoos and Aquaria [32]. The large number of hybrids in our dataset—48 of the 135 true individuals—is also in accordance with captive colony statistics, as 531 of the 1059 individuals in the European Chimpanzee Studbook are hybrids [32].

The impact of narrow sampling in biomedical research outcomes has been repeatedly documented. In mice, where the use of inbred strains is common, the same genotype can lead to very different phenotypes across strains [33,34]; in humans the systematic overrepresentation of individuals of European ancestry has severe implications for the generalisation of medical genetic research and the extent to which its benefits can be translated into other populations [35–37]. Yet, compared to chimpanzees and other primates, humans show a relative dearth of diversity. If not accounting for diversity in human populations can have substantial limitations on the information that can be learnt from them, it is likely that not accounting for the significantly larger amount of variation found across chimpanzee subspecies when doing comparative research is confounding attempts to truly understand humans as great apes.

## Methods

### Sample selection

As of the 19th of January 2021, the NCBI SRA database contained 4389 RNA-seq samples with taxon ID Pan troglodytes, sequenced through 17 different technologies, with the main ones being the Illumina HiSeq 2000 (483 samples), Illumina HiSeq 2500 (2316 samples) and HiSeq 4000 instruments (422 samples). To avoid potential biases from calling SNPs on sequencing data generated with vastly different technologies [38], we considered only samples sequenced on these three instruments, retaining 3221 samples across 49 distinct BioProjects (collections of BioSamples from a single initiative in NCBI) for further consideration. Four single-cell RNA-seq studies [39–42] accounted for 2164 of these samples, which we excluded from further analysis as our genotyping pipeline yields low quality genotype calls from single-cell data due to read scarcity.

The remaining 1059 entries were associated with a total of 808 BioSample IDs; we randomly chose one entry per BioProject for further processing. For studies with more than 5-6 replicates per individual (as identified by metadata on SRA), we only retained 5 randomly chosen replicates in order to reduce computational load; thus our description of sample swaps and mislabelled samples is limited. For paired-ended samples, we retained only the R1 file. Two datasets (PRJNA445737 and PRJDB1766) failed processing with GATK. We additionally excluded 2 datasets (PRJNA385016, PRJNA481380) as the individuals in these studies were already well-represented in our dataset. Our final dataset includes 486 BioSamples, 339 of which were single-ended and 147 pair-ended, from 40 different BioProjects (Supplementary Tables 1 and 2) and spans 20 different tissue or cell types (and 2 samples labelled “pooled tissues”); for simplicity, when plotting we collapsed these tissue/cell types into 9 different categories:

1. Brain: brain, iPSC-derived cortical spheroids, iPSC-derived neural progenitors, iPSC derived neurons
2. Heart: heart, iPSC-derived cardiomyocytes, muscle, myoblast
3. Blood: whole blood, peripheral blood mononuclear cells, lymphoblastoid cell lines, endothelial tissue
4. Skin: skin, dermal fibroblasts
5. iPSCs
6. Kidney
7. Liver
8. Testis
9. Colon

In parallel, we downloaded the complete set of 28,559,256 SNPs genotyped in 59 wild-born chimpanzees from all four subspecies (12 Western, 19 Eastern, 10 Central and 19 Nigeria-Cameroon) generated by De Manuel et al through high coverage whole-genome sequencing [14]. We used CrossMap 0.3.8 [43] to convert genomic coordinates for these SNPs from panTro4 to panTro5 using chain files from the UCSC Genome Browser. 1626 unclassified contigs were then removed with BCFtools 1.9, [44]. Fewer than 1% of SNPs mapped to these contigs.

### SNP Calling

We removed all unclassified contigs from the panTro5 genome release. Then we implemented the GATK 3 version of GATK’s RNAseq short variant discovery (SNPs + Indels) pipeline (https://gatk.broadinstitute.org/hc/en-us/articles/360035531192-RNAseq-short-variant-discovery-SNPs-Indels-). We additionally incorporated an initial quality control step to this process, and used Trimmomatic 0.38 [45] to remove reads with average quality below 20 and minimum length below 40 bp, as some studies had read lengths of 50 bp. We applied Trimmomatic separately on longer samples and ensured that this relaxed filter did not have a sizeable impact on the number of reads kept after trimming.

To determine values for variant calling filters, we considered existing literature [1] and GATK recommendations. Using a test run of 15 samples, we implemented to following filters, which produced enough high quality SNPs for downstream analyses: Fisher Strand (FS) < 26; Total Depth (DP) < 10 (especially relaxed as sequencing depth varies considerably between studies and between genes); Mapping Quality (MQ) > 25; Genotype Quality (GQ) > 20.

### Merging and filtering

A total of 7,529,801 variants passed the variant calling filters above. We then used VCFtools 0.1.15 [46] to remove indels and variants that had failed variant filtering criteria described above, for a final set of 6,943,957 SNPs. We used PLINK 1.90 [47] to calculate a Site Frequency Spectrum for retained SNPs, which suggested widespread presence of duplicate samples (Supplementary Figure 6). As expected, the majority of variants were singletons, which we also removed with VCFtools.

We then used BCFtools to merge our genotype calls with the reference WGS SNPs, and applied different missingness thresholds of 2%, 5% and 10% to both the unmerged and merged datasets. After considering results obtained at these different thresholds (Supplementary Figure 2), we elected to proceed with a missingness threshold of 5%, which allowed us to account for the effect of tissue-specific and temporal expression patterns that characterise the different RNA-seq samples. This resulted in a final set of 54,706 SNPs genotyped in our samples, as well as 11,659 SNPs in our samples and the WGS data.

### Relatedness analyses

We used PLINK to compute genome-wide IBS pairwise distances between all samples. We then subtracted the IBS values from 1 to get a distance matrix. Using the *ape* package (ver 5.4-1, [48]) in R 4.0.5 [49] we then performed hierarchical clustering on these and plotted dendograms using treeio 2.2.4 [50]. As the evolutionary distance between our samples is not consequential, we arbitrarily chose the UPGMA method for dendogram generation. In parallel, we made use of Somalier 0.2.10 [25], which takes polymorphic sites and calculates pairwise relatedness (coefficient of relationship, defined as (shared-hets_i; j_-2*ibs0_i; j_) / (min (hets_i_; hets_j_)) between each pair of samples. Approximately 1000 SNPs with a minor allele frequency as close to 0.5 as possible are needed for a high-confidence analysis; we therefore considered only the 946 SNPs in our data with MAF ≥ 0.25. When filtering replicate or duplicate samples from clusters, we chose at random for sets that contained 2 individuals and retained the one with the highest average relatedness to the rest of individuals in the cluster otherwise.

Since neither set of results provided enough resolution to detect higher-degree relationships, we also used PLINK to calculate pairwise identity-by-descent metrics Z0, Z1 and Z2 to identify relative pairs within our dataset as in [51]. We defined the following thresholds for identifying relatedness between pairs on the basis of known pedigree relationships, metadata, and [51]: parent-offspring: Z1 ≥ 0.7; inferred siblings: Z2 ≥ 0.30, Z1 ≥ 0.30 and Z0 ≤ 0.40; second-degree relatives: 04 ≤ Z1 < 0.7; potential identical: Z2 ≥ 0.65 and Z0 < 0.10. Ternary diagrams were generated with the ggtern3.3.0 [52] package.

#### Population structure and ancestry assignments

We used SmartPCA (part of Eigensoft 7.2.1) [53] to perform PCA in the 59 wild-born samples and confirm that it was possible to recover population structure across the species with low numbers of SNPs. Our results at various missingness thresholds were comparable to those observed when using genome-wide data. We then projected our 486 RNA-seq samples onto this space, again using SmartPCA.

We also used ADMIXTURE 1.3.0 [54] to quantify subspecies ancestry of the RNA-seq samples. We set K = 4, hoping to identify the four chimpanzee subspecies, but the presence of duplicates and significant admixture in the RNA-seq samples (both known and unknown) confounded the algorithm at K = 4 because identical individuals sampled repeatedly clustered together as separate populations. Therefore, we used the WGS samples as reference to perform a supervised ADMIXTURE analysis. RNA-seq samples were assigned a particular subspecies ancestry if that subspecies contributed more than 6.25% to its ancestry, equivalent to an unadmixed great great grandparent.

#### Identification of ambiguous samples

Our initial dendogram of 486 samples included a clade of 18 samples that had consistently large pairwise IBS distances (> 0.1) with all other samples in the data (Supplementary Figure 1). Reasoning that these might represent sample swaps or contamination issues, we mapped and quantified raw reads using Kallisto 0.45.1 [55] against the mitochondrial genomes of chimpanzee (RefSeq NC_001643.1), human (rCRS, NC_012920.1), orang-utan (NC_002083.1) and rhesus macaque (NC_005943.1), as these were the other taxa included in the studies associated with these samples. In all cases, we identified a significant fraction of reads originating outside chimpanzees, confirming our intuition (Supplementary Table 3); other samples from our dataset that fell within the main part of the dendogram did not exhibit this pattern. We therefore removed these samples from all downstream analyses.

### Analysis code

All analyses described were carried out using custom bash and R scripts, and are available at https://gitlab.unimelb.edu.au/navyas/chimpanzee-snp-calling

## Data availability

All datasets in this study are publicly available. A VCF of genotype data from wild-born chimpanzees used as reference is available at https://www.biologiaevolutiva.org/tmarques/data. GEO IDs of all analysed datasets are available in Supplementary Tables 1 and 2. A VCF file containing genotype calls from all 468 RNA-seq samples, unfiltered for missingness, is available for download at Figshare with DOI 10.26188/14822289.

## Acknowledgements

We thank Davis McCarthy, Rob Lanfear, Phillip Bayer and members of the Gallego Romero lab at the University of Melbourne for discussion. We also thank Yoav Gilad, Bryan Pavolvic and Benjamin Fair for sharing a partial pedigree of Yerkes chimpanzees. Michelle Ward confirmed the sample swaps affecting S23; this mislabelling has now been corrected on the SRA record.

## Author contributions

NS and IGR designed the study, performed all analyses and wrote the manuscript. BS implemented the GATK pipeline and provided comments on the manuscript.

## Supplementary Table and Figures

Supplementary information consists of 5 tables and 6 figures.

- **Supplementary Table 1: List of studies**. All studies considered in the sampling process and relevant meta-information.
- **Supplementary Table 2: List of samples**. All chimpanzee-identified samples from 40 studies, including un-analysed replicates. A “Sample_” name was used for every sample during processing.
- **Supplementary Table 3: Mislabelled samples not of chimpanzee origin**. Samples determined to be mislabelled from the results of clustering analysis.
- **Supplementary Table 4: Unexpected relative pairs from Somalier analysis**.
- **Supplementary Table 5: Known and inferred relative pairs from IBD analyses**.
- **Supplementary Figure 1. 486 chimpanzee-identified samples in public RNA-seq data**. UPGMA dendogram was generated from all genotyped samples. 54,706 SNPs were genotyped at ≤ 5% missingness and used to calculate pairwise IBS distances between samples. Non-chimpanzee outlier samples have been indicated, forming a distinct clade from the rest of the dataset.
- **Supplementary figure 2: Principal component analysis of 527 chimpanzee samples with genotype data at different missingness thresholds**. Projection of 468 chimpanzee RNA-seq samples of unknown ancestry onto the PCA of 59 wild-born samples of known ancestry [14] using A. 18,679 SNPs at 10% missingness. B. 11,659 SNPs at 5% missingness. C. 5814 SNPs at 2% missingness. D. 3545 SNPs at 1% missingness. Colors indicate the four distinct chimpanzee subspecies and samples of unknown ancestry from public RNA-seq datasets.
- **Supplementary figure 3: 468 chimpanzee samples in public RNA-seq data**. UPGMA dendogram of 468 samples retained after removing non-chimpanzee outliers. The first column shows the tissue from which the sample was derived. The second column distinguishes different replicate sets from different studies. The particularly extensive clade of cell line PR00818 has been labelled.
- **Supplementary figure 4: Duplicate clusters identified from 237 chimpanzee samples**. UPGMA dendogram of 237 samples, retaining only one replicate per individual in a study. Filled boxes indicate the site from which each individual was sampled (if known). Clusters of potentially duplicate samples, identified from clustering and metadata information, have been labelled numerically, bottom to top. Samples labelled in red are failing to cluster with other duplicates and discussed in the text.
- **Supplementary figure 5: Relationships between non-clustering sample pairs with unexpectedly high Somalier-calculated relatedness**. Ternary diagrams showing pairwise IBD estimates between a subset of 25 samples. Samples were taken from the 18 non-clustering pairs with ≤ 0.5 Somalier-calculated relatedness. All possible pairs including these 25 individuals have been plotted and labelled. Thresholds for defining different relationships between pairs are indicated by black lines on the diagram.
- **Supplementary figure 6: Site frequency spectra in RNA-seq derived genotype data**. **A.** In the full data set B. After removing known replicate samples C. In the full dataset, after filtering for 5% missingness.

