## Supplementary figures and images for "Genetic diversity in chimpanzee transcriptomics does not represent wild populations"

### Supplementary Figure 1

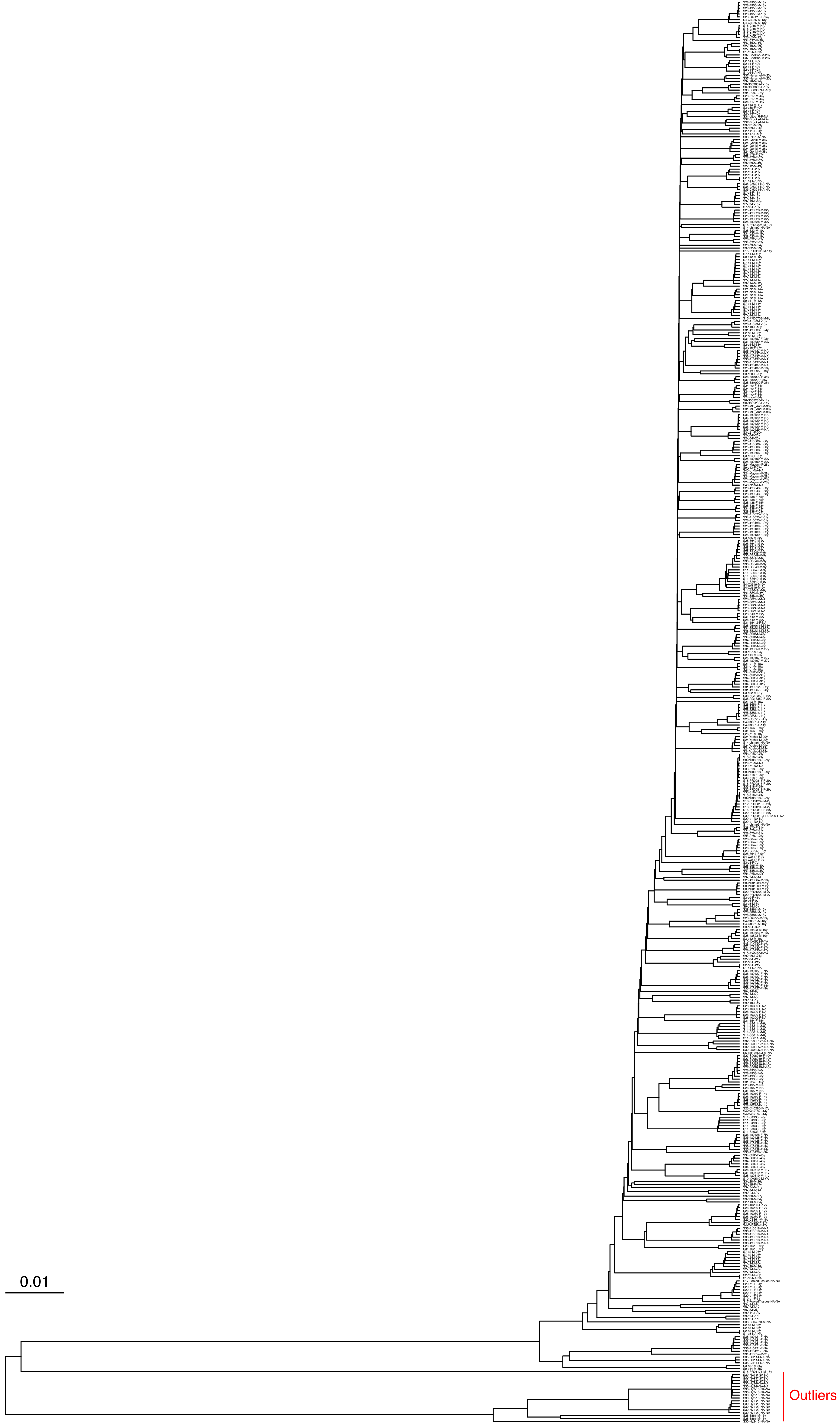

### Supplementary Figure 2

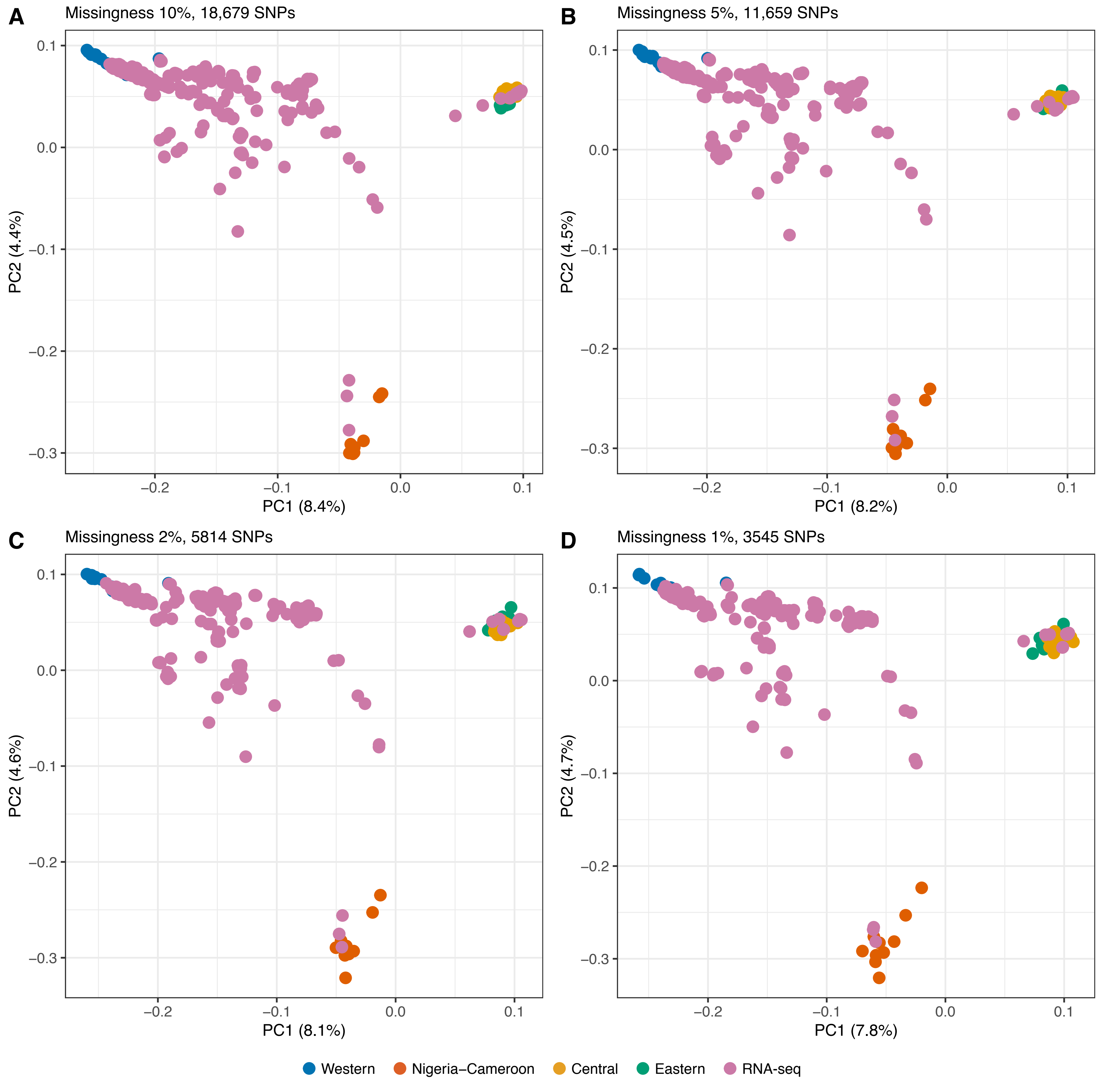

### Supplementary Figure 3

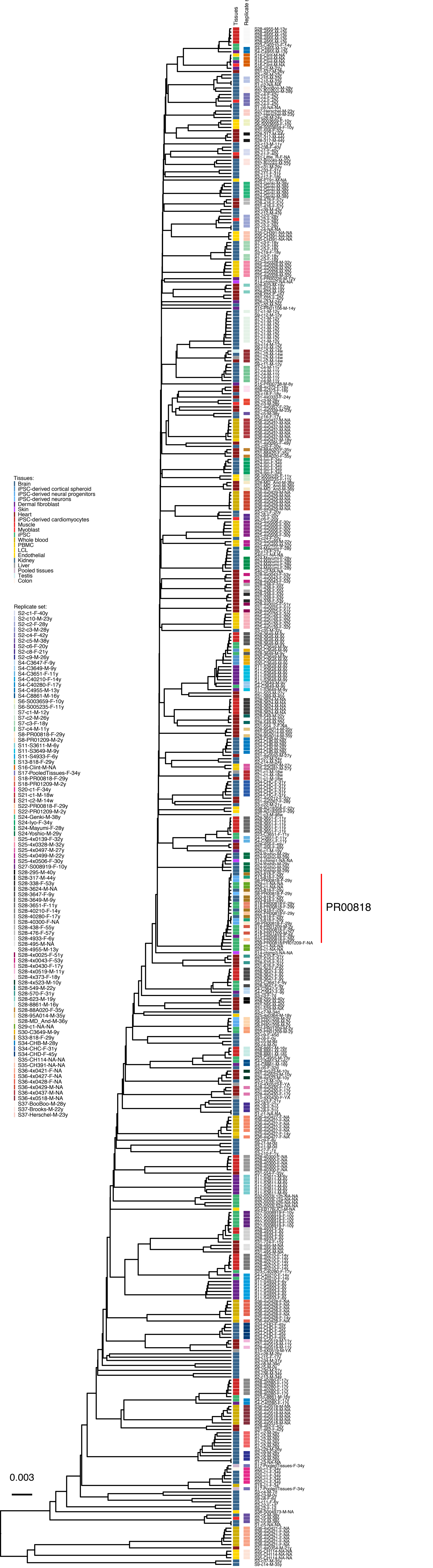

### Supplementary Figure 4

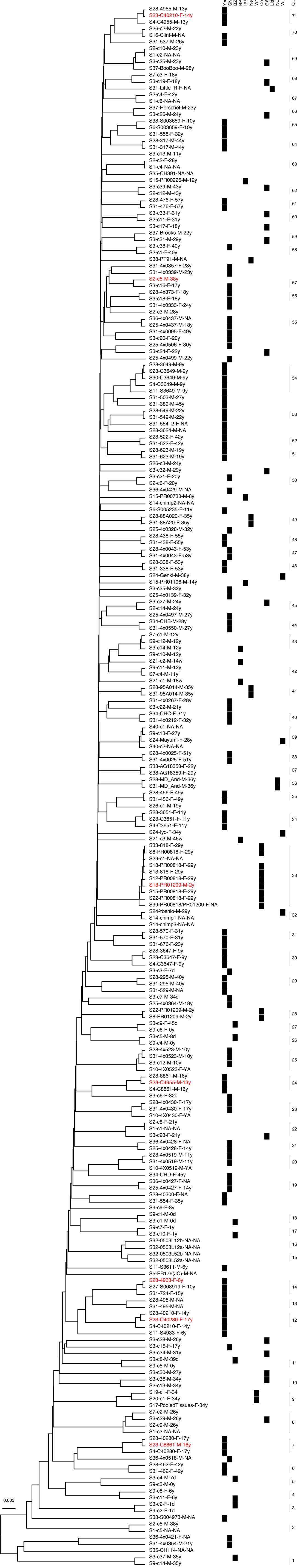

### Supplementary Figure 5

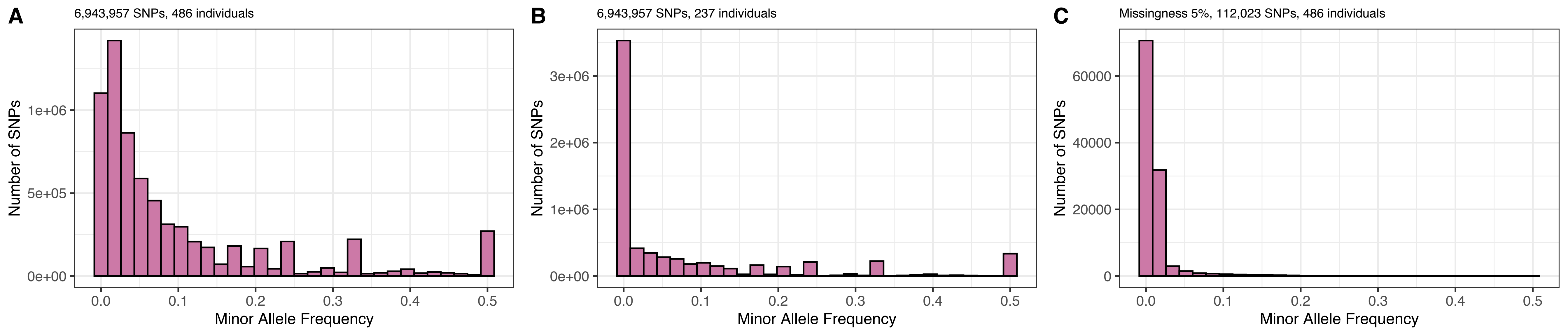

### Supplementary Figure 6

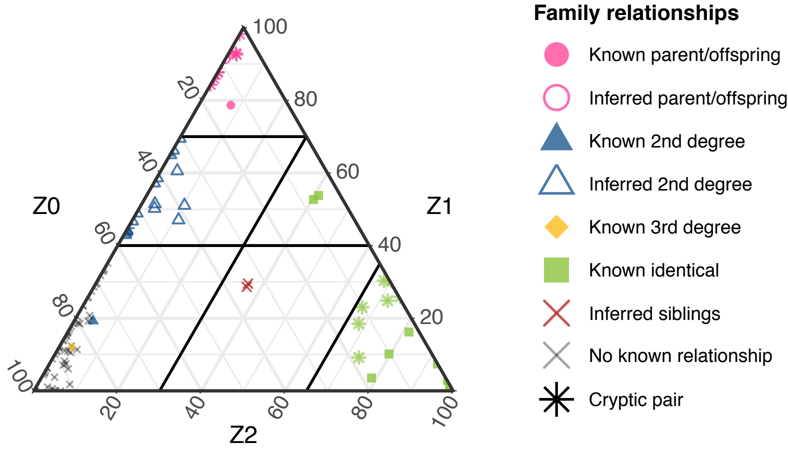
